# *Txn1* mutation causes epilepsy associated with vacuolar degeneration in the midbrain

**DOI:** 10.1101/2021.10.07.463470

**Authors:** Iori Ohmori, Mamoru Ouchida, Hirohiko Imai, Saeko Ishida, Shinya Toyokuni, Tomoji Mashimo

**Affiliations:** Section of Developmental Physiology and Pathology, Education, Institute of Academic and Research, Okayama University, Tsushima 3-chome 1-1, Kita-ku, Okayama 700-8530, Japan; Department of Child Neurology, Medicine, Dentistry and Pharmaceutical Sciences, Okayama University, Shikatacho 2-chome 5-1, Kita-Ku, Okayama 700-8558, Japan; Department of Molecular Oncology, Medicine, Dentistry and Pharmaceutical Sciences, Institute of Academic and Research, Okayama University, Shikatacho 2-Chome 5-1, Kita-Ku, Okayama 700-8558, Japan; Department of Systems Science, Kyoto University Graduate School of Informatics, Yoshida-Honmachi, Sakyo-ward, Kyoto 606-8501, Japan; Division of Animal Genetics, Laboratory Animal Research Center, Institute of Medical Science, The University of Tokyo, Tokyo, 108-8639, Japan; Department of Pathology and Biological Responses, Nagoya University Graduate School of Medicine, 65 Tsurumai-Cho, Showa-Ku, Nagoya 466-8550, Japan

**Keywords:** *Txn1*, Thioredoxin, mitochondria, neuron, vacuolar degeneration, epilepsy, oxidative stress

## Abstract

Thioredoxin (TXN), encoded by *Txn1*, acts as a critical antioxidant in the defense against oxidative stress by regulating the dithiol/disulfide balance of interacting proteins. The role of TXN in the central nervous system (CNS) is largely unknown. A phenotype-driven study of *N*-ethyl-*N*-nitrosourea-mutated rats with running seizures at around five-week of age revealed the relevance of *Txn1* mutations to CNS disorders. Genetic mapping identified *Txn1*-F54L in epileptic rats. The insulin-reducing activity of *Txn1*-F54L rats was approximately one-third that of the wild-type. Vacuolar degeneration in the midbrain, mainly in the thalamus and the inferior colliculus, was observed in the *Txn1*-F54L rats. The lesions displayed neuronal and oligodendrocyte cell death. Neurons in *Txn1*-F54L rats showed morphological changes in the mitochondria. Vacuolar degeneration began at three weeks of age, and spontaneous repair began at seven weeks; a dramatic change from cell death to repair occurred in the midbrain during a restricted period. In conclusion, *Txn1* is essential for the development of the midbrain in juvenile rats.

## Introduction

The brain is more vulnerable to oxidative stress than other organs. Compared to the liver or kidney, the brain has a higher oxygen consumption, in some areas has an increased iron content that catalyzes the generation of reactive oxygen species (ROS), has elevated amounts of lipids with unsaturated fatty acids, and has lower activities of superoxide dismutase (SOD), catalase (CAT), and glutathione peroxidase (GPx) [1]. Several studies have implicated oxidative stress in the development of neurodegenerative diseases such as Alzheimer’s disease, Parkinson’s disease, Huntington’s disease (HD), amyotrophic lateral sclerosis (ALS), and multiple sclerosis (MS) [2, 3, 4].

The primary endogenous sources of ROS in mammals are the mitochondrial respiratory chain and NADPH oxidase (NOX). [5, 6, 7]. Oxidative stress modulates functions ranging from cell homeostasis to cell death [8]. Low levels of mitochondrial ROS production under physiological conditions are required for cellular signaling pathways, such as those regulating proliferation and differentiation. In contrast, excess ROS causes protein denaturation and promotes cell death. The delicate balance between the beneficial and harmful effects of ROS is a vital aspect of living cells and tissues [8].

Living organisms have a defense system that scavenges ROS produced by mitochondria. The enzymatic antioxidant system consists of SOD, CAT, GPx, and thioredoxin (Txn). Txn was first identified as a hydrogen donor for ribonucleotide reductase in *Escherichia coli* [9]. In humans, thioredoxin has been identified in the culture media of adult T-cell leukemia cell lines and Epstein-Barr virus-infected cells [10]. Subsequently, it has been reported that thioredoxin expression is upregulated in many cancer cells and that it inhibits apoptosis of cancer cells [11, 12]. Accordingly, the thioredoxin system has attracted attention as a target for cancer therapy [13, 14, 15]. However, fewer studies have investigated the roles of the TXN system *in vivo.* A previous study reported that transgenic mice overexpressing a dominant-negative mutant of *Txn1* specifically in the heart show cardiac hypertrophy [16]. Homozygous *Txn1* knockout mice are embryonically lethal, whereas heterozygous mice show normal development [17]. Other than these reports, little is known about the effects of reduced TXN activity *in vivo.*

Here, we present a novel animal model for neurodegeneration designated the *Adem* “<u>A</u>ge-<u>de</u>pendent <u>mi</u>tochondrial cytopathy” rat. These animals harbor a *Txn1* missense mutation (F54L). *Adem* exhibit a unique seizure phenotype that appears only at 5 weeks of age. Detailed histological analyses and anatomical analyses using MRI revealed vacuolar degeneration in the midbrain during the epileptic period. Our study reveals a hitherto unknown function of *Txn1* in the CNS.

## Materials and methods

### Animals

We used *Txn1*-F54L rats *(Adem;* Age-dependent mitochondrial cytopathy rat) generated by ENU-mutagenesis and a *Txn1* – T160C (p. F54L) knock-in rat generated by CRISPR-Cas 9. We used the former to characterize the phenotype and the latter to confirm the reproducibility of the phenotype. The *Txn1*-F54L rat was initially discovered in an ENU-induced mutant archive at Kyoto University [18, 19, http://www.anim.med.kyoto-u.ac.jp/nbr] as a rare strain with frequent running seizures only in its juvenile stage. All animal husbandry procedures were performed according to the protocols approved by the institutional experimental animal use committees Okayama University, Osaka University, and Kyoto University. Both sexes were used in the present study.

### Genetic mapping and sequencing analysis of the Adem rat generated by ENU-mutagenesis

The rats with running seizures were backcrossed for more than ten generations on the *F344/NSlc* inbred background (Japan SLC, Hamamatsu, Japan). The repeated backcrosses ensure that the frequency of potential mutations induced by ENU elsewhere in the genome is reduced to approximately 1 in 4×10^6^ bp of the genome. To identify the causative gene, we produced 117 *(Adem×BN/SsNSlc)* × *BN/SsNSlc* backcross progeny. Genetic mapping of the locus was determined in rats that displayed wild running or generalized tonic-clonic seizure phenotypes. Genomic DNA was extracted from rat tail biopsies by an automatic genomic DNA isolation system (PI-200; Kurabo, Osaka, Japan). To localize the locus to a specific chromosomal region, genome-wide scanning on DNA samples was performed using simple sequence length polymorphism (SSLP) markers covering all autosomal chromosomes (Chrs) as previously reported [20].

To enrich the 1.5-Mb region (Chr 5: 75,159,000 to 76,708,000), SureSelect custom DNA probes were designed by SureDesign under moderately stringent conditions, and generated by Agilent Technologies (Santa Clara, CA, USA). The DNA library was prepared using SureSelect reagents and a custom probe kit. Genomic sequence analysis was performed by HiSeq 2500 (2 × 150 bp) according to the standard protocol at Takara Bio.

### Generation of Txn1-T160C (p. F54L) knock-in rat using CRISPR-Cas 9

The possibility of genomic DNA alterations around the *Txn1* gene could not be fully excluded in *Txn1*-F54L rats generated by ENU mutagenesis. As the *Txn1* mutant rats showed a unique phenotype, never previously reported, we generated a new *Txn1*-F54L knock-in rat to test the reproducibility of this phenotype. A *Txn1* T160C (p. F54L) knock-in rat was generated by CRISPR-Cas9 genome editing with long single-stranded DNA (lssDNA), as previously reported [21]. A pair of guide RNAs (gRNAs) targeting intron 1 and intron 2 of rat *Txn1* were designed. LssDNA for *Txn1* was synthesized using a long ssDNA Preparation Kit (Biodynamics Laboratory Inc., Tokyo, Japan) following the manufacturer’s protocol. The sequences of gRNA and ssDNA are listed in Supplementary Table 1. Cas9 mRNA (400 ng/μl), gRNA (200 ng/μl), and ssDNA (40 ng/μl) were delivered into 113 *F344/Jcl* embryos by electroporation as previously reported [22]. The two-cell stage embryos (n = 110) were transferred into the oviducts of five pseudopregnant females. Genomic DNA from founder rats was extracted from the tail biopsies. The CRISPR target site was amplified using the specific primer pairs listed in Supplementary Table 1.

### Genotyping

The last 3 mm of the rat tail was cut at three weeks of age, and DNA was purified with a DNeasy Blood & Tissue kit (Qiagen, Hilden, Germany). A DNA fragment containing a mutation was amplified with a pair of primers as follows: FW 5’ – CCA CAT GGG AGA GTC ACA T −3’ and RV 5’-ATA GCC TGG AAG CGG TCA GAT G −3’ (Sigma-Aldrich Japan, Inc., Tokyo, Japan). The PCR products (521 bp) were then treated with the restriction enzyme *Bsp1286I* (New England Biolabs, Ipswich, MA, USA), and the F54L mutant DNA was digested to 260 bp and 261 bp fragments. Genotyping was determined by the size of the digested fragments on a 3% agarose gel.

### Video monitoring of epileptic seizures

To clarify the appearance and frequency of epileptic seizures, both the heterozygotes (n = 8) and homozygotes (n = 9) were placed under a 24-hour video monitoring system (Handycam HDR-CX480, SONY, Japan) from week three to nine,

### Histological analysis

Brains were fixed with 10% formalin neutral buffer solution. Tissue sections were cut from paraffin-embedded brain samples taken at 2, 3, 5, and 9 weeks of age and were subjected to hematoxylin and eosin (HE) and Bodian and Klüver-Barrera (KB) staining using standard protocols. The stained sections were examined under a BZX-700 microscope (Keyence, Osaka, Japan). We evaluated brain lesions, including samples from the thalamus and the inferior colliculus, in the sagittal and coronal planes.

### Head MRI

To evaluate the extent of brain lesions and their changes over time, an MRI scan was performed in 1 mm slices. Three animals were examined for each genotype at 5, 7, and 9 weeks of age. The same animal was examined twice on days 1 and 14 as the storage period of the MRI-attached breeding room was limited to no longer than two weeks. The mice were anesthetized with isoflurane and laid in the prone position on a cradle. Anesthesia was maintained by inhalation of 2% isoflurane in air at 1.4 L/min through a face mask.

Throughout the MRI measurements, respiratory rate and rectal temperature were monitored using a dedicated system (Model 1025, MR-compatible Small Animal Monitoring and Gating System, SA Instruments, Inc., NY, USA). The body temperature was maintained by a flow of warm air using a heater system (MR-compatible Small Animal Heating System, SA Instruments).

All MR images were obtained with a 4.7 Tesla preclinical MR scanner (BioSpec 47/16 USR, Bruker BioSpin MRI GmbH, Ettlingen, Germany). A quadrature volume resonator (Bruker BioSpin) was used for signal detection. The scanner was operated using ParaVision 6.0.1 (Bruker BioSpin). Two-dimensional multi-slice T2-weighted MR imaging was performed using a rapid acquisition with relaxation enhancement (RARE) sequence. The whole-head images were acquired in three orthogonal (coronal, sagittal, and axial) orientations. The acquisition parameters were as follows: echo time (TE), 12 ms; effective TE, 36 ms; RARE factor, 8; acquisition bandwidth, 62.5 kHz; in-plane spatial resolution, 0.156 × 0.156 mm^2^; slice thickness, 1 mm; slice gaps, 0 mm; the number of averages, 2. For coronal orientation, repetition time (TR), 3000 ms; field of view (FOV), 25 × 29.3 mm^2^, matrix size, 120 × 136 (zero-filled to 160 × 188 before image reconstruction); the number of slices, 24; acquisition time, 1 min 42 s. For sagittal orientation, TR, 2500 ms; FOV, 27 × 27.5 mm^2^, matrix size, 131 × 128 (zero-filled to 173 × 176); number of slices, 18; acquisition time, 1 min 20 s. For axial orientation, TR, 2500 ms; FOV, 27 × 30.5 mm^2^, matrix size, 131 × 144 (zero-filled to 173 × 195); the number of slices, 16; acquisition time, 1 min 30 s.

### Western blotting

Western blotting was used to quantify the different cell types of the brain lesion and Txn1 expression levels in multiple organs in the rats. The protein extracts (10 μg) from rat tissues were mixed with sample buffer solution (Nakalai Tesque, Kyoto, Japan), denatured at 95 °C for 5 min, and loaded onto a sodium dodecyl sulfate (SDS)-polyacrylamide gel (Mini-protean TGX Precast gels; Bio-Rad, Hercules, CA) for electrophoresis. The proteins were blotted onto a polyvinylidene fluoride (PVDF) membrane using the iBlot Dry Blotting System (Thermo Fisher Scientific). Membranes were incubated with anti-Thioredoxin1 rabbit antibody (CST#2298; Cell Signaling Technology, Danvers, MA, USA), anti-NeuN rabbit antibody (ab177487; Abcam, Cambridge, UK), anti-Olig2 rabbit antibody (ab109186; Abcam), anti-glial fibrillary acidic protein (GFAP) rabbit antibody (ab7260; Abcam), anti-Iba1 rabbit antibody (ab178847; Abcam), and anti-glyceraldehyde 3-phosphate dehydrogenase (GAPDH) rabbit antibody (CST #2118). Mouse and rabbit primary antibodies were detected using anti-mouse IgG-HRP-linked antibody (CST #7076) and anti-rabbit IgG-HRP-linked antibody (CST; #7074), respectively. Chemiluminescence was detected using Western Lightning ECL Pro (PerkinElmer Japan, Kanagawa, Japan) and the ChemiDoc Touch (Bio-Rad).

### Immunohistochemistry (IHC) analyses

A standard IHC protocol was used to stain the brain tissue samples. In brief, 5-μm-sized paraffin-embedded tissue sections were de-paraffinized with xylene. Antigen retrieval was performed by autoclaving at 120 °C for 5 min. Endogenous peroxidases were quenched by treatment with 0.3% hydrogen peroxide in methanol for 30 min at room temperature. They were then incubated with blocking serum for 1 h at 22-26 °C. Sections were incubated with the following primary antibodies to determine the cell type in the brain lesion: anti-NeuN rabbit antibody (ab177487; Abcam, Cambridge, UK) and anti-Olig2 rabbit antibody (ab109186; Abcam) for 1 h at room temperature. To determine DNA oxidative injury and lipid peroxidation, sections were stained with anti-8-hydroxy-2’-deoxyguanosine (8-OHdG) mouse antibody (MOG-020P, JaICA, Japan), and anti-4-hydroxynonenal (4-HNE) mouse antibody (MHN-020P, JaICA, Japan) overnight at 4 °C, respectively. The sections were then reacted with a biotinylated rabbit polyclonal secondary antibody (PK-6200, Vector Laboratories, Burlingame, CA, USA). To reveal the staining, we used an avidin-biotinylated peroxidase complex (PK-6200, Vector Laboratories). After washing, the slides were incubated with 3,3’-diaminobenzidine (DAB) (SK-4100, Vector Laboratories) and immediately washed with tap water after color development. The slides were then counterstained with hematoxylin, mounted with dibutyl phthalate xylene (DPX) and observed under a BZX-700 multifunctional microscope (Keyence, Osaka, Japan). Quantification of IHC was performed by assessing five randomly selected fields in the inferior colliculus and thalamus. The percentages of positive cells or positive areas were calculated using BZX Analyzer software (Keyence).

### Transmission electron microscopy (TEM)

Samples were immersed in 2% glutaraldehyde and 2% paraformaldehyde in 100 mM phosphate buffer (PB) for 24 h at 4 °C. They were then postfixed with 2% osmium tetroxide in 100 mM PB for 1.5 h at 4 °C, after which they were rinsed with 100 mM PB, and dehydrated through a graded series of ethanol treatments. They were embedded in Spurr resin (Polysciences Inc., Warrington, PA, USA), cut into ultrathin sections, and stained with uranyl acetate and lead citrate. The ultrathin sections were observed using a Hitachi H-7650 TEM (Hitachi High-Tech Corp., Tokyo, Japan).

### Plasmids and purification of recombinant proteins

For recombinant Thiredoxin1 proteins, rat *Txn1* cDNAs were amplified by PCR using LA-Taq polymerase (Takara Bio Inc, Shiga, Japan) with FW-primer: 5’-GGA TCC ATG GTG AAG CTG ATC GAG AG-3’; and RV-primer, 5’-GTC GAC TTA GCT GTC CAT GTG CTG GCG TTC GAA TTT AGC GGT TTC TTT GAA TTC GGC AAA CTC CGT AAT AGT GG-3’ (Sigma-Aldrich Japan, Inc., Tokyo, Japan). The PCR products were cloned into a pCMV-Tag2 plasmid (Stratagene, San Diego, CA, USA) digested with *BamHI* and *SalI* containing FLAG-tag and S-tag sequences. The DNA sequence of the plasmids was confirmed by DNA sequencing using a BigDye Terminator FS Ready-Reaction Kit (Applied Biosciences, Little Chalfont, Buckinghamshire, UK) and an ABI 3130x Genetic Analyzer (Applied Biosciences). The plasmids were transfected into HEK293 human embryonic kidney cells using Lipofectamine2000 (Invitrogen, Carlsbad, CA, USA). The recombinant proteins were purified with anti-FLAG agarose affinity gel (A2220; Sigma-Aldrich Japan), eluted with FLAG peptide (A3290; Sigma-Aldrich Japan), and filtered using an ultrafiltration membrane (Amicon Ultra 3 K; Millipore, Burlington, MA, USA).

### Insulin-reducing activity assay

Thioredoxin activity was measured using a thioredoxin activity assay kit (Redoxica, Little Rock, AR, USA), according to the manufacturer’s instructions. In brief, the purified recombinant thiredoxin1 proteins or protein extracts from rat thalamus and cortex were reacted with insulin and NADPH in assay buffer, and thioredoxin activity was determined as a change in the amount of oxidized NADPH by measuring the decrease in absorbance at 340 nm per minute using a spectrophotometer DeNovix DS-11 (DeNovix, Tokyo, Japan).

### Primary fibroblast and neuron culture

Newborn rats were sacrificed under deep anesthesia with isoflurane on the day of birth. The animals were dissected in the surgical suite and disinfected with 70% alcohol. For primary fibroblast culture, the abdominal skin tissues were cut into small pieces in DMEM (Fujifilm Wako Pure Chemical Corporation, Osaka, Japan) and disrupted by pipetting. The cell suspension was transferred and maintained in DMEM supplemented with 10% fetal bovine serum (Thermo Fisher Scientific, Waltham, MA, USA), 500U/ml penicillin, and 500 ug/ml streptomycin (Sigma-Aldrich Japan, Inc., Tokyo, Japan) in a fully humidified atmosphere of 5% CO_2_. After about two weeks, fibroblast cells were cultured in DMEM supplemented with 10% fetal bovine serum, 100U/ml penicillin, and 100 ug/ml streptomycin.

For primary neuronal culture, we referred to a protocol using a postnatal mouse [23] and modified it. We used the cerebral cortex of rats on the first postnatal day. Tissues were treated with 0.125% trypsin (Thermo Fisher Scientific) and 0.004% DNase-I (Sigma-Aldrich) at 37 °C for 15 min and dissociated mechanically. Cells were plated on poly-l-lysine and laminin-coated glass-bottomed 35-mm culture dishes. Cells were maintained in a culture medium with DMEM, 100 μg/ml penicillin-streptomycin (Thermo Fisher Scientific) and 10% fetal calf serum (Thermo Fisher Scientific) at 37 °C in a humidified incubator with 95% air and 5% CO_2_. On the second day, the culture medium was replaced with a medium containing DMEM, 2% B-27 supplement (Thermo Fisher Scientific), and 5% fetal calf serum. The culture medium was refed with a medium containing DMEM, α-MEM, F-12 nutrient mixture, 2% B-27 supplement, 0.34% glucose, 25 μM 5-fluoro-deoxyuridine, 25 μM uridine, one mM kynurenic acid (Sigma-Aldrich), and 1% fetal calf serum on the fourth day. Cultures were used for the experiments on days 9–15. Mitochondria and nuclei were stained using MitoBright LT Red (Dojindo, Kumamoto, Japan) and Cellstain-Hoechst33258 solution (Dojindo, Kumamoto, Japan), respectively.

### Terminal deoxynucleotidyl transferase (TdT) dUTP Nick-End Labeling (TUNEL) assay

For determining apoptosis, paraffin-embedded brain tissues at three weeks of age and primary fibroblasts were assessed using the In Situ Cell Death Detection Kit, TMR red (Sigma-Aldrich Japan, Inc., Tokyo, Japan), following the manufacturer’s instructions. We compared cell death during incubation of primary fibroblasts with and without the addition of 300 μM hydrogen peroxide for 3h. In brief, the cells were cultured on an 8-chamber culture slide (BD Falcon, Franklin Lakes, NJ, USA), washed with phosphate-buffered saline (PBS), fixed with 10% formalin neutral buffer solution (Fujifilm Wako Pure Chemical Corporation, Osaka, Japan) for one h at room temperature, and treated with permeabilization solution for 2 min on ice. TMR red-labeled dUTP was incorporated into 3’-OH DNA ends by terminal deoxynucleotidyl transferase, and the fluorescence signal was detected using a BZX-700 microscope (Keyence, Osaka, Japan) or a fluorescence microscope (Olympus IX71; Olympus Corporation, Tokyo, Japan).

### Statistical analysis

Results are expressed as mean ± SEM. The sample size for each experiment is indicated in the figure and figure legends. One-way analysis of variance (ANOVA) was used to compare between three groups. Statistical significance was set at p < 0.05. Bonferroni adjustment was used to determine the statistical differences between the groups. All significant statistical results are indicated within the figures using the following conventions: * p < 0.05, ** p < 0.01, *** p < 0.001.

## Results

### Discovery of a novel epileptic rat with Txn1 missense mutation

A strain exhibiting unique running seizures was identified in an archive of ENU-mutated rats. Thus, we decided to identify the responsible gene. By analyzing simple sequence length polymorphism (SSLP) markers on mutant (n = 21) and sibling (n = 25) DNA pools, the *Adem* locus was roughly mapped on chromosomes 2 and 5. The locus was subsequently refined by 26 and 33 SSLP markers for each chromosome, and mapped to a 1.5-Mb genomic region, between markers D5Mit17 (Chr 5: 75,159,412-75,159,546) and D5Rat113 (Chr 5:76,707,365-76,707,563) (RGSC_v3.4) (Fig. 1A). Ten known or predicted genes within the *Adem* region *(Ptpn3, Palm2, AC134204.1, Akap2, LOC685849, Txn1, Txndc8, Svep1, Musk, Lpar1)* were identified from the Ensembl database (http://www.ensembl.org).

**Fig. 1.**
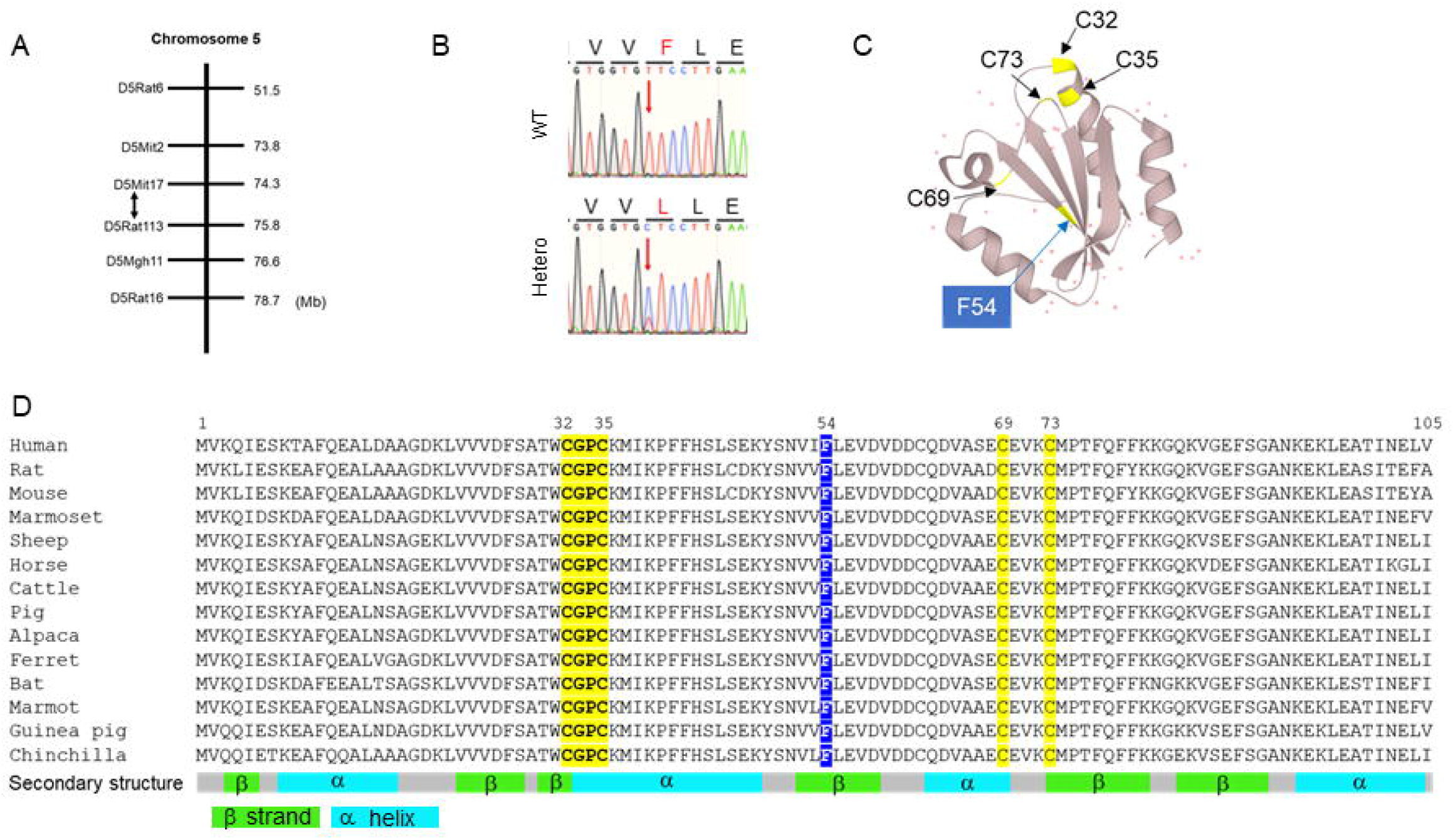
Discovery of a novel epileptic rat with a *Txn1* missense mutation. (A) *Adem* locus mapped to a 1.5-Mb genomic region, between markers D5Mit17 and D5Rat113. (B) DNA sequencing exhibiting c. T160C in exon 3 of the *Txn1* gene, causing a substitution of phenylalanine for leucine at residue 54. (C) Functional sites and F54 were plotted in the 3D structure of human thioredoxin. (D) Alignments of the sequence of Txn1 gene in mammals.

We performed next-generation sequencing analysis of genomic DNA from 75,159,000 to 76,708,000 on Chr5 using the SureSelect target enrichment approach (Agilent). In mutant DNA, we found only one heterozygous missense mutation in the exon region, c. T160C, in exon 3 of the *Txn1* gene (NM_053800, ENSRNOG00000012081.6) by comparing the parental strain F344/NSlc. This variant caused a substitution of phenylalanine for leucine at residue 54 (Fig. 1B).

Txn1 is widespread in prokaryotes and eukaryotes, and its amino acid sequence is well conserved in mammals (Fig. 1D). Amino acid sequences are 90% identical in humans and rats. Since the structure of rat thioredoxin has not been reported, we plotted the position of F54 on the 3D structure of human thioredoxin (UniProt, https://www.uniprot.org/). The positions of amino acids with important functions in Txn1 were plotted on the structure. Txn1 has an active center sequence Cys32-Gly-Pro-Cys35 motif and carries out redox reactions by dithiol/disulfide exchange reactions between Cys32 and Cys35 [24, 25, 26]. Both Cys69 and Cys73 are nitrosylated in response to nitric oxide (NO) [26, 27]. Cys-73 can serve as a donor for the nitrosylation of target proteins. The location of the F54L mutation is remote from these active sites (Fig. 1C and Fig. 1D), suggesting that it is not directly involved in the dithiol/disulfide exchange reaction or S-nitrosylation. However, the PolyPhen-2 prediction (http://genetics.bwh.harvard.edu/pph2/index.shtml) found that the F54L mutation is possibly damaging with a score of 0.785, and the Sorting Intolerant From Tolerant (SIFT, http://www.ngrl.org.uk/Manchester/page/sift-sorting-intolerant-tolerant.html) sequence is deleterious.

### Vacuolar degeneration appears during an epileptic period

Video monitoring showed that the frequency of seizures peaks at 5-week of age in both heterozygotes (Fig. 2A) and homozygotes (Fig. 2B). There were no differences in seizure frequencies between heterozygotes and homozygotes.

**Fig. 2.**
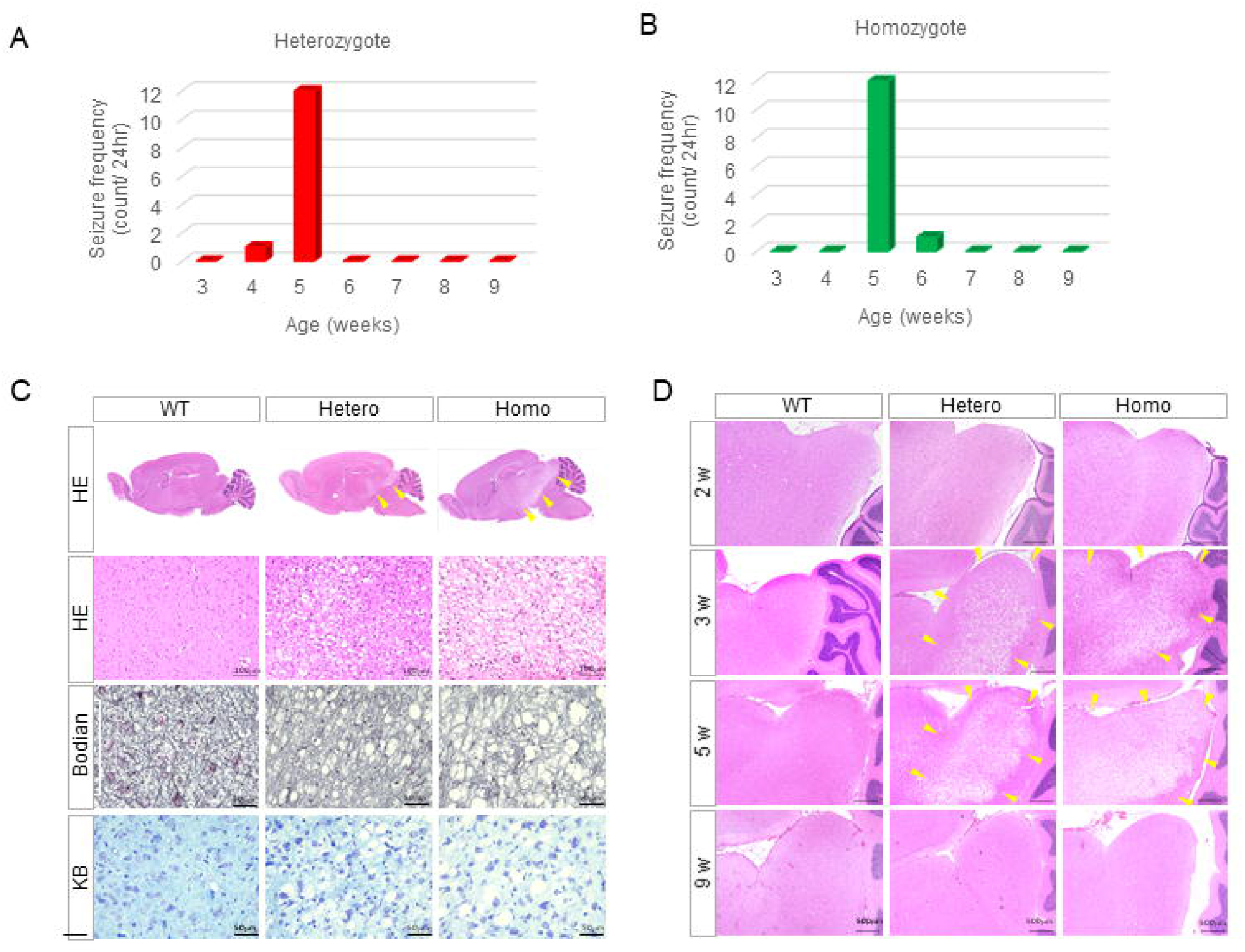
Vacuolar degeneration appears during an epileptic period. (A) Total seizure frequency of heterozygotes at each week (N = 8). (B) Total seizure frequency of homozygotes at each week (N = 9). (C) Histology at five weeks of age. At least three animals of each genotype were examined. (D) The representative changes of inferior colliculus lesions from 2 to 9 weeks of age for each genotype.

Next, a histological examination was performed at five weeks to determine whether a structural change was the cause of epilepsy. Surprisingly, vacuolar degeneration was found in the inferior colliculus and thalamus of the mutants. (Fig. 2C). No changes were observed in the hippocampus, which is prone to epilepsy. Bodian staining, which is used to observe nerve fibers, and Klüver-Barrera (KB) staining, which is used to observe the myelin sheath, exhibited decreased staining in the mutants. These histological tests showed that homozygotes had more severe lesions than heterozygotes. We examined when these lesions had started developing (Fig. 2D). There were no remarkable changes in the inferior colliculus at two weeks of age. However, vacuoles became evident in the inferior colliculus and thalamus at three weeks of age and were then widespread at five weeks of age. They almost disappeared completely after nine weeks.

### MRI showed transient high signals of T2-weighted images in the midbrain

MRI detected hyperintense regions of T2-weighted images (T2WI) spread around the thalamus in mutant rats at five weeks of age (Fig. 3A). The hyperintense regions of the heterozygous and homozygous T2WI were symmetrical. The lesions were localized in the thalamus, inferior colliculus, superior colliculus, and hypothalamus in homozygotes (Fig. 3B). The hyperintensity region of T2WI in homozygous rats was wider than that in heterozygous rats (see also Supplementary Fig. S1, S2, and S3). Temporal lesions were also examined using MRI in the same individual. Serial MRI examination revealed that the lesions at three weeks spread or were similar in size at five weeks. However, the lesions shrank at seven weeks and disappeared at nine weeks (Fig. 3C). The weekly changes observed on MRI were similar to those seen in the histological examination.

**Fig. 3.**
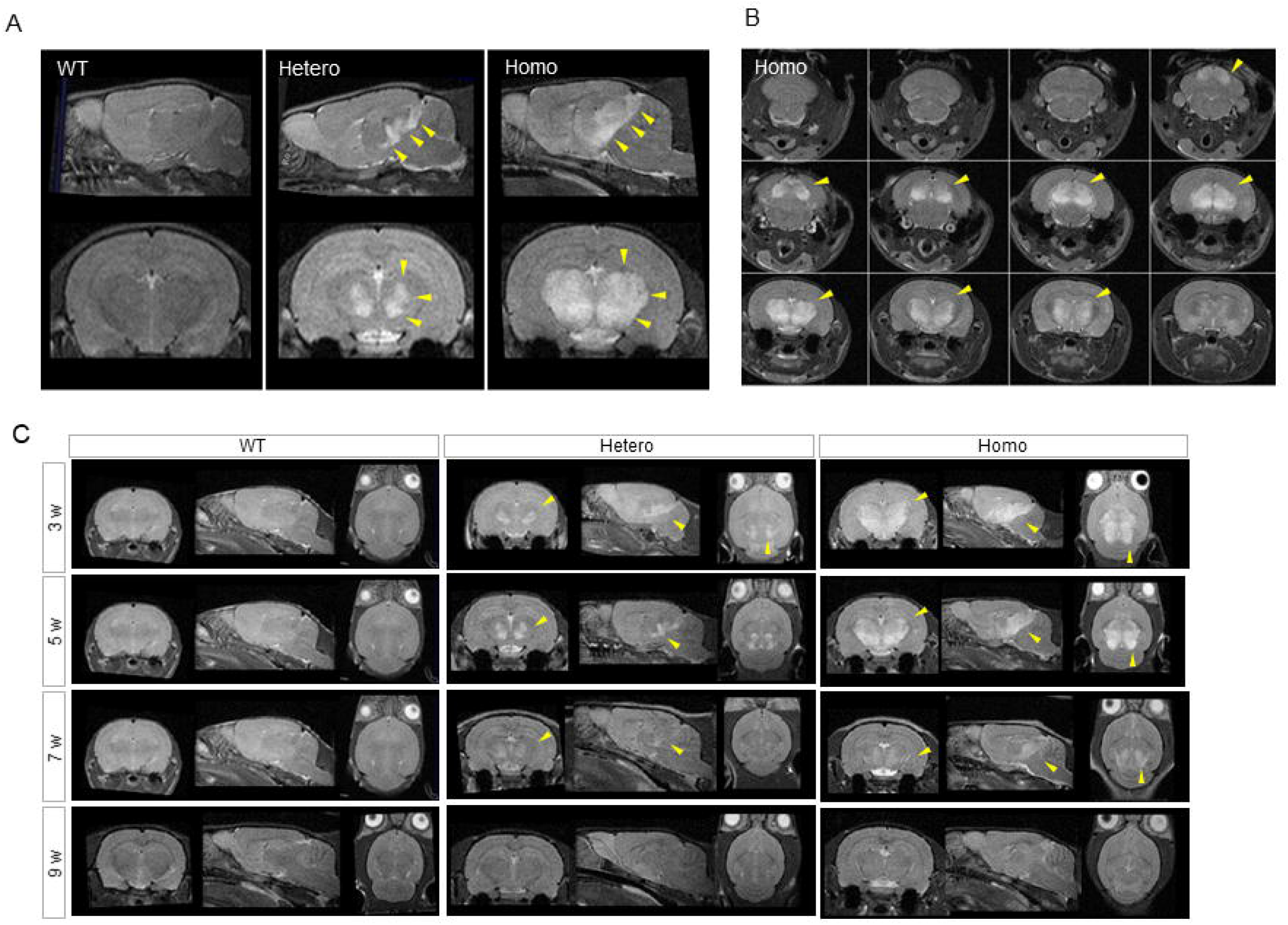
MRI demonstrates transient high signals of T2-weighted images in the midbrain. (A) The representative MRI findings at five weeks for each genotype (sagittal slices). (B) The coronal slices of homozygotes at five weeks. (C) The temporal changes of the brain lesion for each genotype from three to nine weeks.

### Neuronal and oligodendrocyte cell loss occurs in the midbrain

Western blotting of the midbrain using NeuN, Olig2, GFAP, and anti-IbaI antibodies showed a decrease in NeuN and Olig2 and an increase in GFAP and IbaI expression in heterozygotes and homozygotes (Fig. 4A). Immunohistochemical studies showed significantly decreased numbers of NeuN-positive neurons (Fig. 4B and 4C) and Olig2-positive oligodendrocytes (Fig. 4D and 4E). NeuN-positive neurons in heterozygotes and homozygotes had larger cell bodies than wild-type (WT) rats. Electron microscopy of the lesions in the mutant rats revealed that microtubules in the vacuoles and myelin sheath were wrapped around the vacuoles (Fig. 4H and 4I). These findings indicate that the vacuoles were dilated axons of the neurons. Examination of the mitochondrial morphology showed sparse cristae in the mutant, even though the neurons retained their structures (Fig. 4J and 4K).

**Fig. 4.**
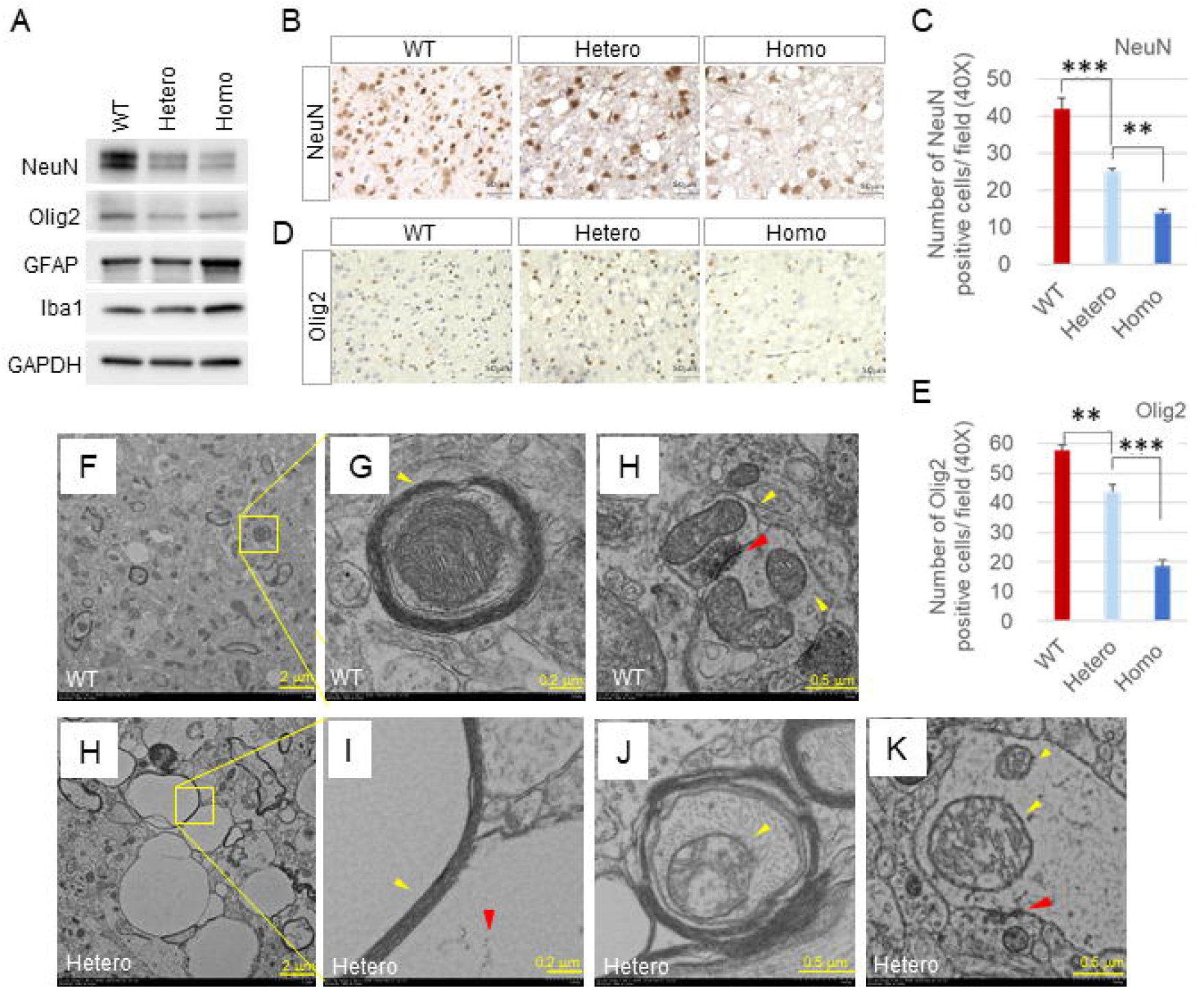
Neuronal and oligodendrocyte cell loss occurs in the midbrain. (A) Protein levels of each cell in the midbrain at five weeks. Anti-NeuN, anti-Olig2, anti-GFAP, and Iba1 bodies are markers of neurons, oligodendrocytes, astrocytes, and microglia. (B) IHC of the anti-NeuN-positive cells in the inferior colliculus for each genotype. (C) Quantification of NeuN-positive cells. Quantification was performed by assessing five randomly selected fields in the inferior colliculus and thalamus for each genotype. (D) IHC of Olig2-positive cells in the inferior colliculus for each genotype. (E) Quantification of Olig2-positive cells. Quantification was performed by assessing five randomly selected fields in the inferior colliculus and thalamus for each genotype. (F) Representative TEM image in the inferior colliculus of WT rat at four weeks. (G) Magnification of the axon in (F). The yellow arrowhead indicates myelin sheath. (H) Representative TEM image of the synapse in the inferior colliculus of WT rat at four weeks. The yellow arrowhead indicates mitochondria with normal morphology, and the red arrowhead indicates postsynaptic density, corresponding with the synaptic terminal. (I) Magnification of the vacuole in (H). The yellow arrowhead indicates the myelin sheath, and the red arrowhead indicates a microtubule. (J) Representative TEM image of the axon in the midbrain of the heterozygous rat at four weeks. The yellow arrowhead indicates mitochondria with sparse cristae. (K) Representative TEM image of the synapse in the inferior colliculus of WT rat at four weeks. The yellow arrowhead indicates mitochondria with sparse cristae, and the red arrowhead indicates postsynaptic density.

### Txn1-F54L shows decreased insulin-reducing activity

Fig. 5A shows the thioredoxin/peroxiredoxin system. Four central enzyme systems defend against oxidative stress by scavenging ROS generated in the body. These include SOD, which scavenges superoxide, CAT, GPx, and thioredoxin-dependent peroxiredoxin (Prdx).

**Fig. 5.**
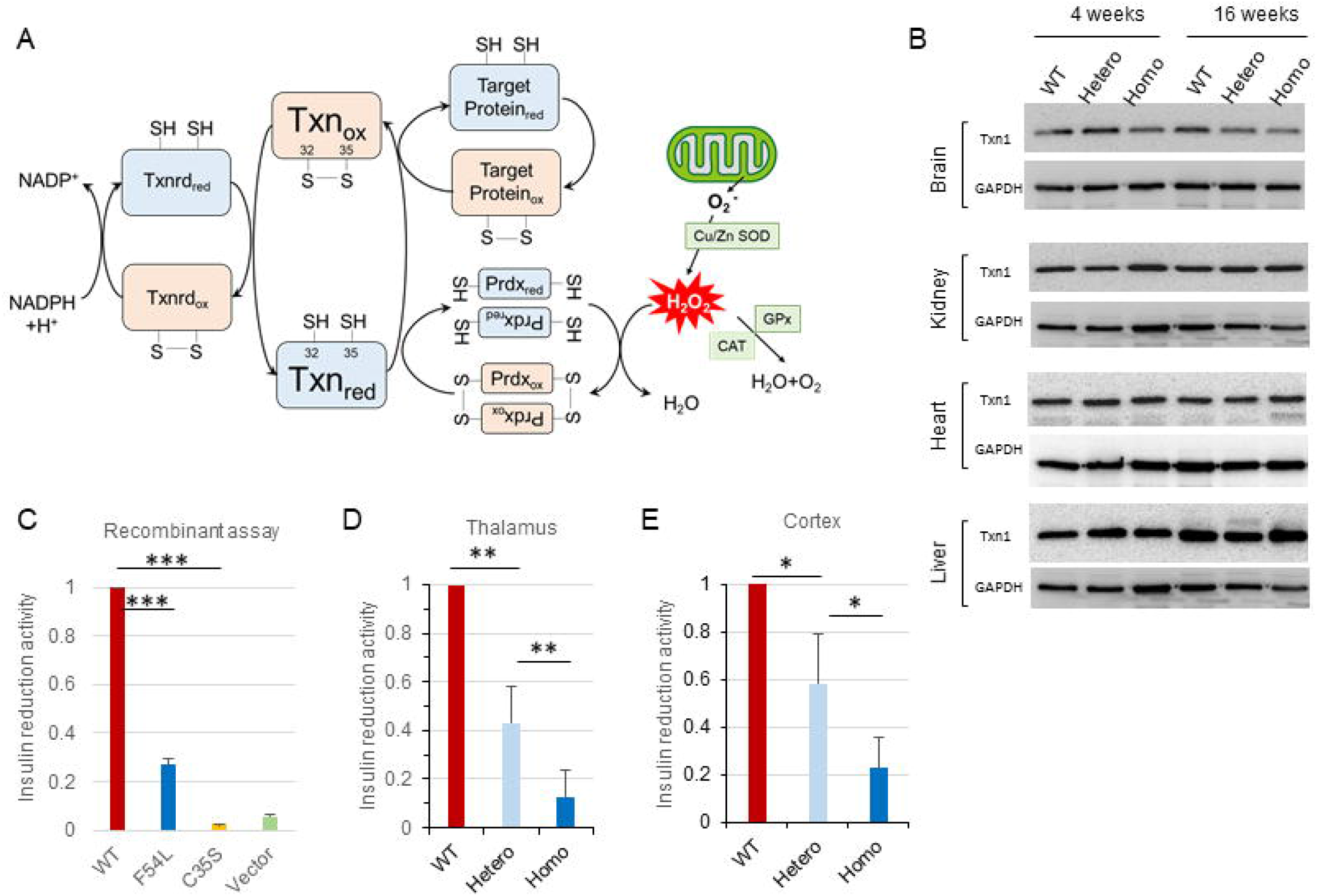
*Txn1*-F54L shows decreased insulin-reducing activity. (A) Thioredoxin/peroxiredoxin system. (B) Protein level of thioredoxin in the brain, kidney, heart, and liver. (C) Insulin-reducing activity determined using recombinant proteins. (D) Insulin-reducing activity of the thalamus for each genotype. (E) Insulin-reducing activity of the cortex for each genotype.

Thioredoxin forms a system with nicotinamide adenine dinucleotide phosphate (NADPH) and thioredoxin reductase (Txnrd) to reduce disulfide bonds in target proteins. Thioredoxin is responsible for converting the target protein from oxidized to the reduced form and scavenging H_2_O_2_ via Prdx. To confirm whether the reduced expression of Txn1 underlies the changes in the brain during 3–5 weeks of age, we examined the expression level of Txn1 protein in the brain, kidney, heart, and liver by western blotting at four weeks of age when vacuoles appeared and at 16 weeks of age when repair occurred. We found that Txn1 was equally expressed in all organs, and there was no change between 4 and 16 weeks of age (Fig. 5B).

Next, we measured thioredoxin activity by using recombinant protein (Fig. 5C), the thalamus at five weeks of age, where vacuolar degeneration was observed (Fig. 5D), and the frontal cortex (Fig. 5E), where vacuolar degeneration did not occur. Our results showed that the insulin-reducing activity of *Txn1*-F54L was decreased to approximately one-third of that of the WT. Similarly, insulin-reduction activity in the thalamus was reduced in heterozygotes and reduced much further in homozygotes compared to that in WT mice (Fig. 5D).

Interestingly, reduced activity was also observed in the cortex, where there was no vacuolar degeneration (Fig. 5E).

### Oxidative damage in the midbrain and susceptibility to cell death under oxidative stress

Next, we investigated the pathogenesis underlying the vacuolar degeneration. Staining with 8-OHdG, which indicates oxidative stress-induced DNA damage, and 4-HNE, which indicates lipid oxidation, showed brain staining in the mutant lesions (Fig. 6A). The area of 8-OHdG (Fig. 6B) and 4-HNE (Fig. 6C) positive cells were significantly increased in heterozygous and homozygous rats compared to WT cells. TUNEL assay of paraffin-embedded sections, including the thalamus, showed slight staining in heterozygous and homozygous rats at three weeks of age (Fig. 6D). Next, we examined cellular vulnerability to H_2_O_2_ using primary fibroblasts and cortical neurons derived from WT and homozygous rats. The TUNEL assay showed that homozygous fibroblasts were slightly stained even under standard culture medium, and the staining became significantly stronger when H_2_O_2_ was added. In contrast, TUNEL-positive cells in WT fibroblasts did not increase with the addition of 0.3 mM H_2_O_2_ (Fig. 6E). We also found that WT primary neurons showed no significant changes, whereas homozygous neurons showed cell death and nuclear rupture on treatment with 0.3 mM H_2_O_2_ (Fig. 6F).

**Fig. 6.**
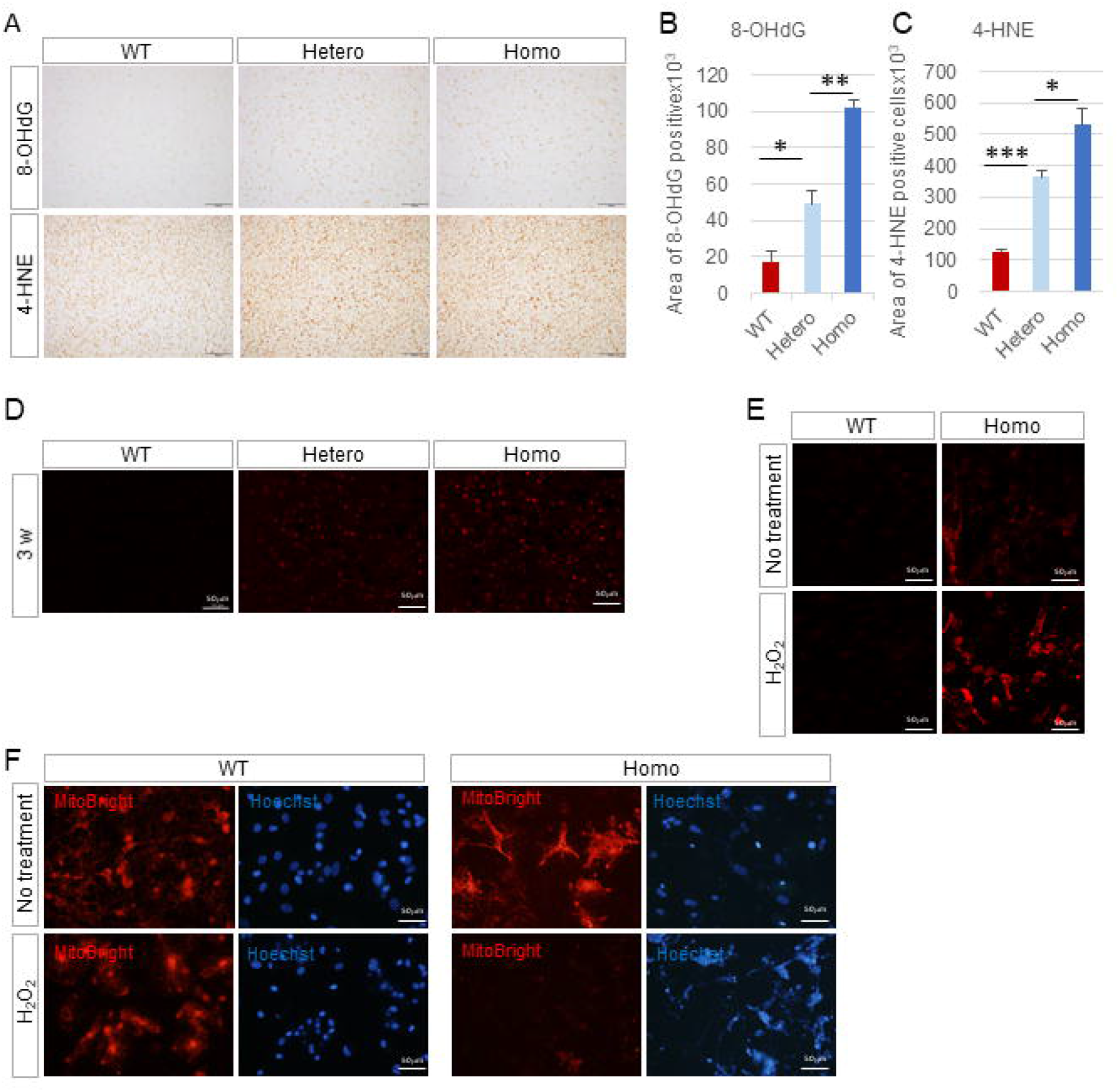
Oxidative damage in the midbrain and susceptibility to cell death under oxidative stress. (A) Oxidative damage in the midbrain was assessed by IHC using anti-8-OHdG antibody and 4-HNE. (B) Quantification of 8-OHdG-positive cells for each genotype. (C) Quantification of 4-HNE-positive cells for each genotype. (D) TUNEL assay of the paraffin-embedded sections including thalamus for each genotype. (E) TUNEL assay of the primary fibroblasts derived from the WT and homozygous rats. TUNEL staining was performed under the standard medium with or without 0.3 mM H_2_O_2_. (F) Mitochondria and nucleus of primary neurons derived from WT and homozygous rats treated with standard medium with or without 0.3 mM H_2_O_2_.

### Txn1-F54L rat generated by genome editing replicates vacuolar degeneration in the midbrain

Finally, we conducted reproducibility experiments to confirm whether the *Txn1*-F54L mutation is the sole cause of the F344/NSlc phenotype. Genome editing using CRISPR-Cas9 was used to generate F344/Jcl rats with the *Txn1-*F54L mutation. We placed 5-week-old F344/Jcl *Txn1*-F54L heterozygous rats under 24-hour video monitoring and recorded running seizures eight times/24 h (a total of N = 5) (Supplementary Video 2). Running seizures and transition from running to tonic seizures were observed. These seizure symptoms were similar to those observed in *Adem* rats harboring *Txn1*-F54L. The lesions were localized in the thalamus, hypothalamus, and superior and inferior colliculi and were more extensive in homozygous than in heterozygous rats (Fig. 7A and 7B). The vacuoles in the midbrain started developing at two weeks of age, became apparent at three weeks, and recovered at nine weeks (Supplementary figure 4). Although the genetic background of F344/NSlc generated by ENU-mutagenesis and F344/Jcl is somewhat different, transient vacuolar degeneration localized in the midbrain was reproduced in F344/Jcl rats carrying the *Txn1*-F54L mutation.

**Fig. 7.**
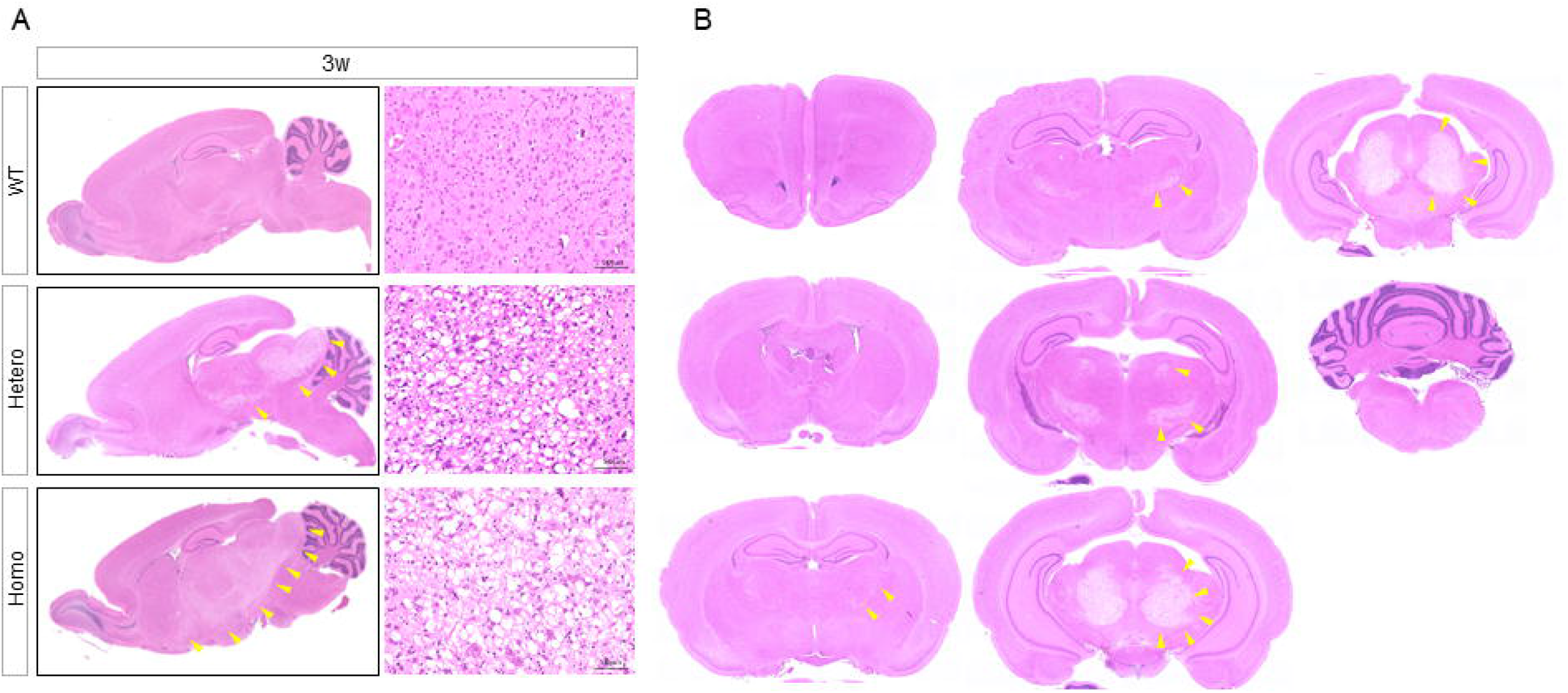
*Txn1*-F54L rat generated by genome editing replicates the vacuolar degeneration in the midbrain. The yellow arrowheads indicate vacuolar degeneration. (A) Representative HE staining of the sagittal brain section at three weeks for each genotype. (B) The coronal slices of the brain for each 500 mm section in the heterozygote.

## Discussion

The function of TXN in vivo remains unclear. Txn1 is located in the cytoplasm, nucleus, and extracellular region [25, 28], whereas Txn2 is located in the mitochondria [29]. A homozygous *TXN2* mutation has been linked to an infantile-onset neurodegenerative disorder with severe cerebellar and optic atrophy and peripheral neuropathy [30]. To the best of our knowledge, there are no reports of *TXN1* mutations that cause human diseases. Herein, we discovered a link between *Txn1* mutations and CNS disorders in mammals for the first time. The loss of Txn1 function leads to neuronal death with abnormal mitochondrial structure in the midbrain.

Txnrd catalyzes the reduction of oxidized Txn in an NADPH-dependent manner. Genetically modified mice with inactivation of Txnrd1 or Txnrd2 show embryonic lethality [31, 32], so the Txn/ Txnrd systems have not been elucidated in vivo beyond the fact that they are essential for fetal development. Other important interacting proteins include the Prdx family, which scavenges H_2_O_2_. Knockout mice for *Prdx1, Prdx2, Prdx3, Prdx4,* or *Prdx5* exhibited various kinds of disorders such as hemolytic anemia, metabolic abnormalities, inflammation, cancer, and age-related phenotypes [33]. The discovery of *Adem* rats may shed light on new roles of the Txn/ Txnrd systems and Txn/ Prdx systems.

*Adem* rats exhibited running seizures and a transition from running to tonic seizures. The focus on this unique epileptic seizure has led to a significant discovery. Running seizures are rarely reported as symptoms of epileptic seizures. According to reports from the 1980s and the 1990s, electrical stimulation of the inferior colliculus and midbrain reticular formation [34, 35, 36] or intense acoustic stimulation [37] induced running seizures. In addition, some evoked running seizures are followed by tonic seizures [35, 36, 37]. The seizure symptoms in the *Adem* rats share similarities with these symptoms. The time of spontaneous appearance and suppression of epileptic seizures and the time of onset and repair of the midbrain lesion were almost the same. This suggests that the cause of epilepsy is a midbrain lesion involving the inferior colliculus and thalamus.

Upon observing the unique vacuolar degeneration in the midbrain of *Adem* rats, we had several questions. First, is there a reduction in cell number in the midbrain? Second, is there an alteration in the reducing activity of *Txn1*-F54L? Third, are the cells derived from *Adem* rats more prone to cell death in response to excess ROS? Fourth, why are vacuoles confined to the midbrain at 3–5 weeks of age? We have discussed our findings based on these questions below.

Western blotting and immunohistochemistry showed a reduction in both neurons and oligodendrocytes in the *Adem* rats. The cell bodies of NeuN-positive cells were more significant than in the WT, and pronounced axon swelling was identified by TEM. These findings suggest that vacuoles are involved in neuronal and oligodendrocyte cell death. Neuronal activity controls oligodendrocyte development and myelination [38, 39, 40], and neuronal cell death inhibits myelination. Further, as neurons and oligodendrocytes are closely related during development, cell death can occur.

Our recombinant protein assay showed that the insulin-reducing activity in the mutant rats decreased to approximately one-third that of the WT. In addition, reduced enzymatic activity was observed in the thalamus, where there was a lesion, and in the cortex, where no lesion was present. F54L is located far from the CGPC motif, which plays a central role in TXN activity. However, PolyPhen-2 and SIFT predicted that the F54L mutation was a loss-of-function mutation. The missense mutation may cause a structural change in the region where the dithiol/disulfide exchange reaction occurs.

In primary cultured neurons and fibroblasts derived from *Adem* rats, cell death was induced more readily than in the WT in the culture medium with H_2_O_2_. IHC with 8-OHdG and 4-HNE, and TUNEL assays stained more intensely in the brain lesions in *Adem* rats than in the WT. These results suggest that the cells in *the Adem* rats are vulnerable to excessive ROS. Txn1 has anti-apoptotic functions in various cells [41, 42, 43]. Loss of function of Txn1 can cause cell death [44, 45, 46]. Proteins that are associated with apoptosis through interaction with Txn1 include apoptosis signal-regulating kinase 1 (ASK1) [47, 48] and Caspase-3 (CASP3) [49]. In addition to apoptosis, other mechanisms of neuronal death could be involved, such as phagocytosis, pyroptosis, and ferroptosis [50]. Western blotting of 5-week-old brains showed a dense band reacting to anti-Iba1 antibody, suggesting an increase in microglia. Microglial activation is a common feature of neurodegenerative diseases. Activated microglia have been reported to kill neurons by releasing TNF-α, glutamate, reactive oxygen, and nitrogen species, which can cause apoptosis, excitotoxicity, and necrotic death of surrounding neurons [51, 52].

Our results could not clearly determine why the midbrain vacuoles were confined to 3–5 weeks of age. The *Txn1* gene is ubiquitously expressed throughout the body *(TXN* thioredoxin [Homo sapiens (human)] – Gene – NCBI https://www.ncbi.nlm.nih.gov/gene/7295). Western blotting of *Adem* rats also showed that thioredoxin was expressed in several organs other than the brain. The insulin-reducing activity of thioredoxin was reduced to one-third of that of the WT, both in the thalamus, where vacuolar degeneration appeared, and in the cerebral cortex, where vacuolar degeneration did not appear. Furthermore, *Adem* rat-derived primary cultured fibroblasts and neurons showed a similar level of cell death induced by excessive ROS. These results of *in vitro* experiments indicate that cell death can be caused by excessive ROS, regardless of the cell type, under the same cell culture conditions. The curious phenomenon of limited onset of neuronal death in the midbrain may be due to the location- and age-specific microenvironment of the midbrain. For example, the neurodevelopmental stage at 3–5 weeks of age might trigger an increased mitochondrial ROS leakage due to high glucose metabolism in the midbrain. The F54L mutation-specific inability of *Txn1* to reduce essential proteins for maintaining ROS homeostasis in the midbrain or the dependence of the midbrain region on the thioredoxin/peroxiredoxin antioxidant system might be high. The thioredoxin/peroxiredoxin system, and not the glutathione system, was reported to be a significant contributor to mitochondrial H_2_O_2_ removal in the brain [53].

Another surprising phenomenon in *Adem* rats is that repair of the vacuolar lesions can already be seen at seven weeks on MRI, and they are almost entirely repaired at nine weeks. Both astrocytes and microglia have been reported to be involved in cytotoxicity and tissue repair. Astrocytes have long been thought to inhibit neuronal repair, but it has been noted that glial activation induces neurogenesis and acts in repair [54, 55]. Microglia phagocytose and remove debris and dead neurons, and release BDNF and IGF to repair damage to neurons [56, 57, 58]. In vacuolar lesions, glial cells may play roles in repair and impart neuropathic effects, or their roles may change from damage to repair. Although this is an exciting point, it is far from the purpose of this study, which was to identify a causative gene of epilepsy and clarify the underlying pathology. We will study this in the future.

We have not yet identified the precise functions disturbed by Txn1-F54L because Txn1 interacts with a wide variety of proteins [59, 60, 61]. This is a limitation of this study. Oxidative stress is responsible for many neurodegenerative diseases [2, 3, 4] and cerebral ischemic-reperfusion injury [62, 63]. The vacuolar degeneration in *Adem* rats may share a common pathological pathway with these neurological disorders. If a common pathway is identified, *Adem* rats would be an attractive animal model. New therapeutic agents for oxidative stress-related diseases can be evaluated more easily because neuronal death occurs spontaneously in a short time and does not require surgical treatment.

## Conclusions

We have shown that the *Txn1*-F54L mutation confers epilepsy. The underlying pathology of *Adem* rats with *Txn1*-F54L is the loss of neurons and oligodendrocytes in the midbrain. *Txn1*-F54L had a reduced insulin-reducing activity compared to that of the WT. Primary cultured cells, both fibroblasts and neurons, derived from *Adem* rats exhibited cell death under excess H_2_O_2_. The mutation might be affecting the midbrain microenvironment in a specific manner leading to the death of neurons and oligodendrocytes in that region. Further studies are required to determine this midbrain-specific function of Txn1.

## List of abbreviations

CNS: central nervous system;
WT: wild type;
ROS: reactive oxygen species
NADPH: Nicotinamide adenine dinucleotide phosphate;
NOX: NADPH oxidases;
ENU: N-ethyl-N-nitrosourea
SOD: superoxide dismutase;
CAT: catalase;
HD: Huntington’s disease;
ALS: amyotrophic lateral sclerosis;
MS: multiple sclerosis;
GSH: Glutathione;
GPx: glutathione peroxidase;
Prdx: peroxiredoxin
Txn: thioredoxin;
MRI: magnetic resonance imaging;
8-OHdG: 8-hydroxy-2’-deoxyguanosine;
4-HNE: 4-hydroxynonenal
TUNEL: Terminal deoxynucleotidyl transferase-mediated dUTP Nick End Labeling
PBS: phosphate-buffered saline;
HE: hematoxylin and eosin;
KB: Klüver-Barrera
DAB: 3,3’-diaminobenzidine;
IHC: immunohistochemistry;
GFAP: glial fibrillary acidic protein;
GAPDH: Glyceraldehyde 3-phosphate dehydrogenase
TEM: Transmission electron microscope;

## Author contributions

T.M discovered the *Adem* rat; I.O. and M.O. performed the phenotypic characterization of this strain; T.M. and S.I. created the genome-edited mutant rats; H.I. supervised the MRI examination and analyzed the data; S.T. supervised the histopathological experiments; Conceptualization, I.O., M.O., and T. M.; Investigation, I.O., M.O., H.I., S.I., S.T., and T.M.; Writing – Original Draft, I.O.; Writing – Review & Editing, M.O. H.I., S.I., S.T., and T. M.; Project Administration, I.O.; Funding Acquisition, M.O., I.O., and T.M. All authors approved the final manuscript.

## Acknowledgments

We thank Ms. Yumiko Morishita, Mika Monobe, and Miki Kajino for technical assistance with histology and IHC. We thank Ms. Masumi Hurutani for the technical assistance with EM. This work was supported by Grants-in-Aid for Scientific Research (16H05354), Grant-in-Aid for epilepsy research from the Japan Epilepsy Research Foundation, and JSPS KAKENHI Grant Number JP16H06276 (AdAMS).

We would like to thank Editage (www.editage.com) for English language editing.

---

**Supplementary Fig. 1.** Representative MRI for WT at five weeks.

(A) Sagittal slices.

(B) Coronal slices.

(C) Horizontal slices.

**Supplementary Fig. 2.** Representative MRI for heterozygote at five weeks.

(A) Sagittal slices.

(B) Coronal slices.

(C) Horizontal slices.

**Supplementary Fig. 3.** Representative MRI for homozygote at five weeks.

(A) Sagittal slices.

(B) Horizontal slices.

**Supplementary Fig. 4.** *Txn1*-F54L rat generated by genome editing replicates the transient vacuolar degeneration in the midbrain from two to nine weeks of age.

