## Supplementary figures and images for "*Txn1* mutation causes epilepsy associated with vacuolar degeneration in the midbrain"

### Supplementary figure 1

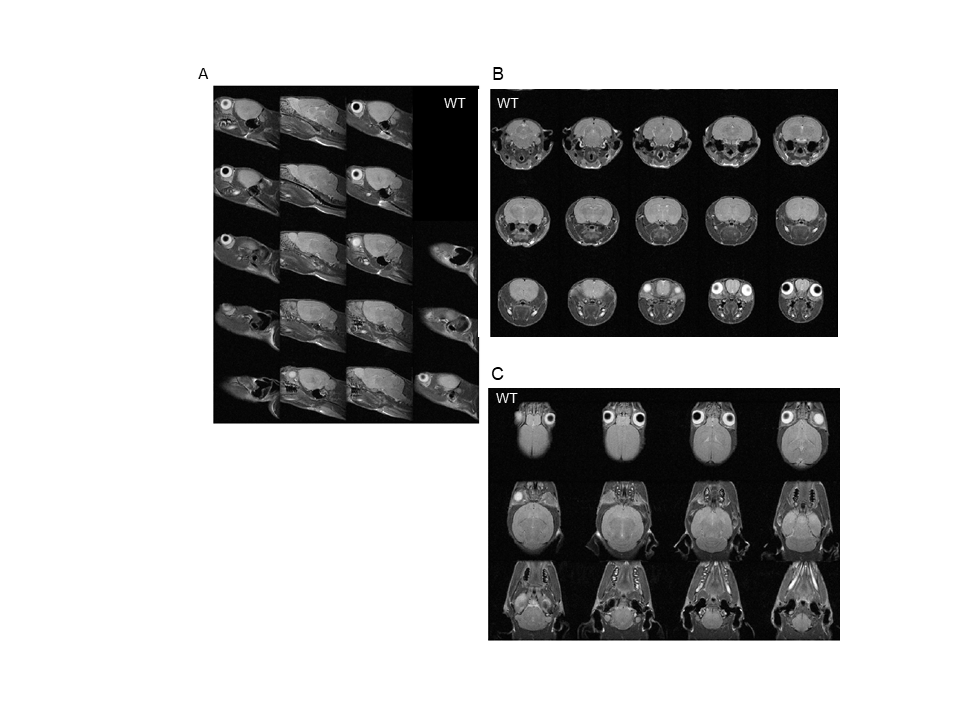

### Supplementary figure 2

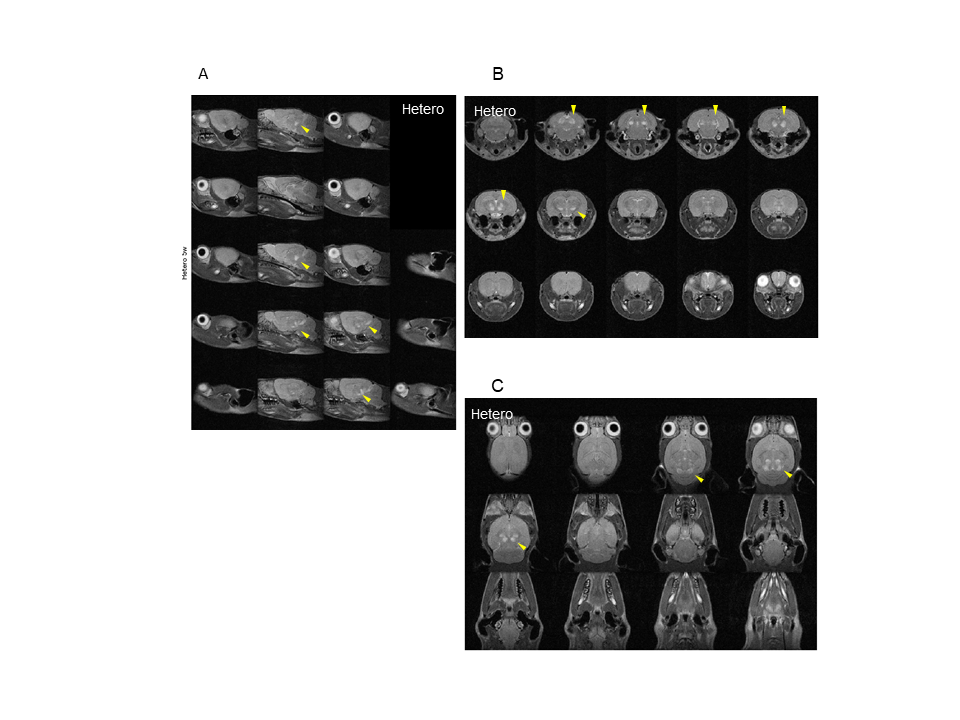

### Supplementary figure 3

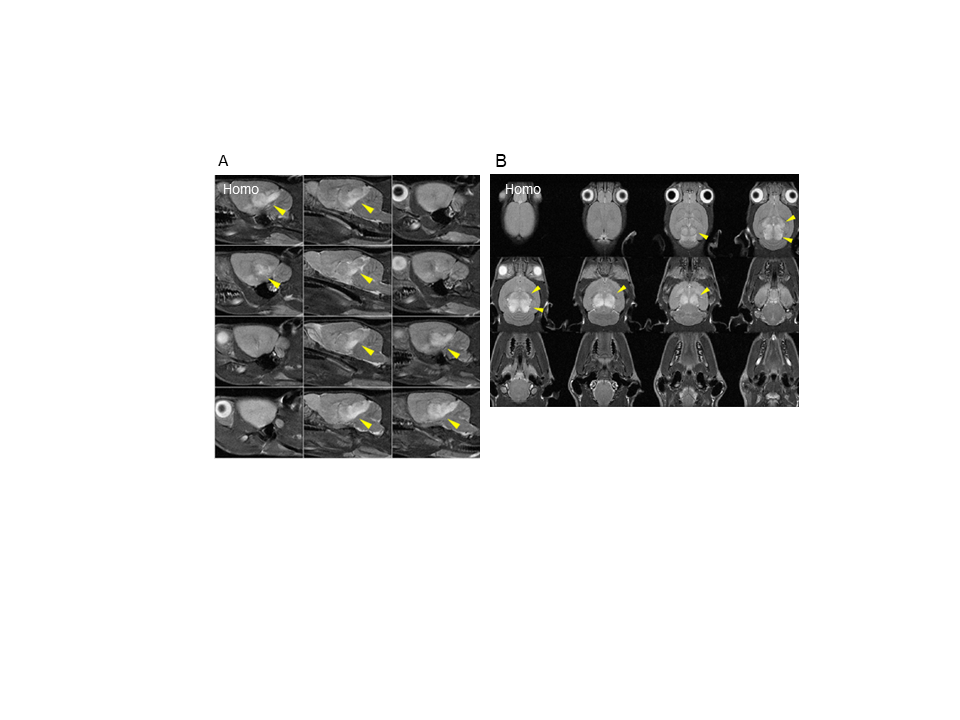

### Supplementary figure 4

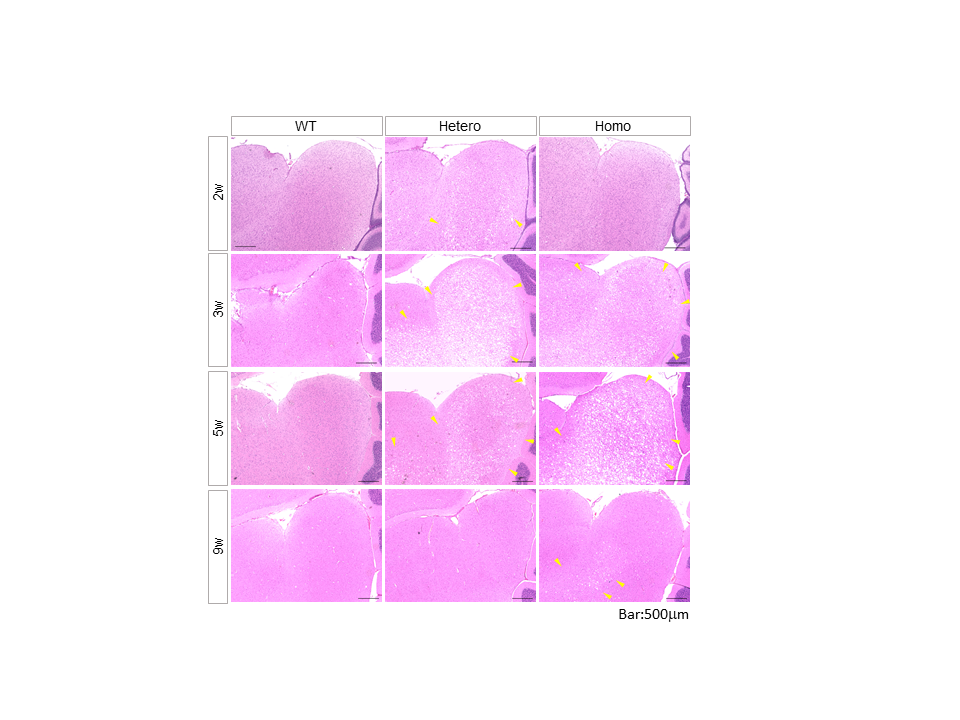
